# Deterioration of Glutaraldehyde Crosslinked Heterograft Biomaterials due to Advanced Glycation End Product Formation and Serum Albumin Infiltration

**DOI:** 10.1101/2020.05.28.117739

**Authors:** Christopher A. Rock, Samuel Keeney, Andrey Zakharchenko, Hajime Takano, David A. Spiegel, Abba M. Krieger, Giovanni Ferrari, Robert J. Levy

**Author notes:** Contact for corresponding author: Robert J. Levy, MD, Division of Cardiology, Department of Pediatrics, The Children’s Hospital of Philadelphia, 3615 Civic Center Blvd, Philadelphia, PA 19104.

## Abstract

Bioprosthetic heart valves (BHV) are fabricated from glutaraldehyde cross-linked heterograft tissue, such as bovine pericardium (BP) or porcine aortic valves. BHV develop structural valve degeneration (SVD), often with calcification, requiring BHV replacement. Advanced glycation end products (AGE) are post-translational, non-enzymatic carbohydrate protein modifications. AGE are present in SVD-BHV clinical explants and not detectable in unimplanted BHV. Here, we studied the hypothesis that BHV susceptibility to AGE formation and serum protein infiltration results in deterioration of both leaflet collagen structure and mechanical properties. In vitro experiments studied BP and porcine collagen sponges (CS) for susceptibility to AGE formation using ^14^C-glucose and ^14^C-glyoxal with and without bovine serum albumin (BSA), as a model serum protein. The results showed AGE formation is a rapid and progressive process. BSA co-incubations reduced glyoxal and glucose uptake by BP and CS. Incubating BP in BSA caused a substantial increase in BP mass, enhanced by glyoxal co-incubation. Per two-photon microscopy, BP with AGE formation and BSA infiltration each induced significant disruption in collagen microarchitecture, with loss of collagen alignment and crimp. These effects are cumulative with the greatest disruption occurring when there was both AGE formation and BSA infiltration. Uniaxial testing of CS demonstrated that AGE formation, together with BSA uptake compared to controls, caused a significant deterioration in mechanical properties with a loss of viscoelastic relaxation and increased stiffness. It is concluded that AGE-BSA associated collagen structural disruption and deterioration of mechanical properties contribute to SVD.

## 1. INTRODUCTION

Heart valve disease is a worldwide problem, affecting millions[1]. At present, there is no effective medical therapy. Patients requiring treatment must undergo either attempted repair of the diseased valves or replacement with a prosthesis. The current preferred replacement valves in adults are bioprosthetic heart valves (BHV) fabricated from glutaraldehyde-fixed xenografts, fabricated from either bovine pericardium (BP) or porcine aortic valves[1]. BHV are relatively non-thrombogenic compared to mechanical heart valve prostheses; this reduces the need for anti-coagulants[2]. However, BHV functional lifespans are limited due to the progressive development of structural valve degeneration (SVD), most often involving leaflet calcification[3,4]. Much of the prior research concerned with mitigating SVD focused on the inhibition of calcification as the primary mechanism[5–9]. However, this yielded only incremental improvements in BHV lifespan[10–13]. Prior studies of explanted BHV showed that the degree of calcification does not completely explain SVD, and that approximately 25% of failed valves lack any significant calcification[14].

The present studies investigated the hypothesis that advanced glycation end products (AGE) contribute to the pathophysiology of SVD. AGE result from the post-translational, non-enzymatic modification of proteins by hexoses or hexose breakdown products from Amadori reactions or other oxidative related mechanisms[15–17]. AGE both modify proteins involved in normal physiologic functions, such as serum albumin and hemoglobin (HA1C) and are associated with the pathophysiology of a number of important diseases including diabetes[18,19] and Alzheimer’s Disease[20–22]. The importance of AGE for SVD is incompletely understood. A prior study by our group evaluated a clinical cohort of 45 explanted BHV with SVD and demonstrated via immunohistochemistry (IHC) that AGE and human serum albumin were present in all explants but were undetectable in unimplanted BHV[23]. This BHV explant study was complemented by *ex vivo* pulse duplicator experiments demonstrating that exposure to both glyoxal, a common AGE intermediary, and serum albumin significantly impaired the hydrodynamic performance of trileaflet, clinical grade BHV[23].

The present studies investigated the hypothesis that BHV are highly susceptible to AGE modification due to both glucose and glyoxal addition and serum albumin uptake, and that these events disrupt the structure of BHV leaflets, adversely affecting biomechanical performance. To test this hypothesis, experiments were performed to determine the following: the glycation kinetics of BHV tissues, the effects of glutaraldehyde-fixation on glycation, the impact of serum albumin exposure on glycation, glycation’s capacity to alter BHV collagen microarchitecture, and glycation’s capacity to alter glutaraldehyde-fixed collagen’s linear elastic and viscoelastic properties.

## 2. MATERIALS AND METHODS

### 2.1 Materials

Bovine serum albumin (BSA, protease-free, >98% purity), D-glucose, glyoxal, sodium azide, sodium chloride, sodium borohydride, and HEPES were purchased from Sigma-Aldrich (St. Louis, MO). Fresh BP were shipped on ice from Animal Technologies (Tyler, TX). Surgifoam® hemostatic collagen sponges (CS) composed of gelatin purified from porcine skin were purchased from Ethicon (Somerville, NJ). The ^14^C-radiolabeled D-glucose (5mCi/mmol) and glyoxal (110mCi/mmol) were purchased from American Radiolabeled Chemicals (St. Louis, MO). Biosol and Bioscint were purchased from National Diagnostics (Atlanta, GA). All other sources of materials have been indicated in individual methods.

### 2.2 Glutaraldehyde Fixation of Bovine Pericardium and Collagen Sponges

BP in a fresh state, shipped on ice, were rinsed in saline (0.9% NaCl) and any residual fatty or muscular tissue was removed by dissection. The BP and CS were immersed for 7 days at room temperature 0.6% glutaraldehyde (Polysciences Inc.; Warrington, PA) in HEPES buffer solution (50mM HEPES, 0.9% NaCl, pH 7.4). The samples were then rinsed in fresh HEPES buffer solution for 1 hour before transferring to storage solution of 0.2% glutaraldehyde in HEPES buffer solution and stored at 4°C. Prior to any experiments, samples were exhaustively rinsed with phosphate-buffered saline (PBS) to remove the storage solution.

### 2.3 Radiolabeled Glycation Assays

#### 2.3.1 Glycation of Glutaraldehyde Fixed Collagenous Tissues

To quantify the glycation kinetics of the model substrates, samples (n=10 for BP and n=5 for CS) were punched from glutaraldehyde-fixed BP and CS using an 8mm biopsy punch. These samples were incubated for 1, 3, 7, 14, or 28 days at 37°C shaking at 110RPM in solutions (1ml per sample) of PBS with either: glucose (100mM), glucose (100mM) + BSA (5%), glyoxal (50mM), or glyoxal (50mM) + BSA (5%). All incubations contained sodium azide (0.1%) to maintain sterility. The glucose media contained ^14^C-labeled glucose (0.544μCi/ml). The glyoxal media contained ^14^C-labeled glyoxal (0.356μCi/ml). The radioactivity of each medium was measured using a Beckman LS 6000 (Beckman Coulter; Brea, CA). Following incubations, the samples were extensively rinsed with deionized water and lyophilized for at least 48 hours. The dried samples were weighed and then digested using Biosol and combined with Bioscint for scintillation counting. The scintillation count was used to calculate glucose or glyoxal incorporation into the substrate normalized by dry weight.

#### 2.3.2 Glycation Comparison of Fresh and Fixed Bovine Pericardium

To compare the glycation capacity of fresh BP relative to glutaraldehyde-fixed BP, samples (n=5) were punched from fresh BP and glutaraldehyde-fixed BP using an 8mm biopsy punch. Samples were rinsed with PBS and then incubated for 28 days in PBS with either: glucose (100mM), glucose (100mM) + BSA (5%), glyoxal (50mM), or glyoxal (50mM) + BSA (5%). Sterility was maintained by adding sodium azide (0.1%) to each incubation. Prior to incubation, radiolabeled reagents were added as above. To calibrate the radioactive signal and confirm sufficient excess of radioactive reagents, a 500μl aliquot was taken from each incubation medium at the start and end of incubation, and radioactivity levels determined. To measure each aliquot’s radioactivity, each aliquot had 5ml of Bioscint added, was shaken till the solution turned clear, and then underwent scintillation counting. At the conclusion of the incubations, samples were exhaustively rinsed with deionized water and underwent a minimum of 48 hours of lyophilization. The lyophilized samples were weighed and then digested in Biosol as above.

#### 2.3.3 Glycation Kinetics of Bovine Serum Albumin

To study the glycation kinetics of albumin *in vitro*, BSA (5%) was incubated with either glucose (100mM) or glyoxal (50mM) for 28 days at 37°C shaking at 110 RPM. The solutions contained either ^14^C-labeled glucose (0.544μCi/ml) or ^14^C-labeled glyoxal (0.356μCi/ml) as above. To maintain the sterility of reaction, the mixture was passed through a 0.2μm syringe filter and sodium azide (0.02%) was added. At 1 day, 3 days, 7 days, 14 days, and 28 days samples were taken from the reacting mixture and passed through Zeba Spin size-exclusion centrifuge columns (ThermoFisher; Waltham, MA) to separate glycated BSA from unbound glucose or glyoxal. To estimate the degree of glycation, each of triplicate samples was mixed with Biosol and measured using a liquid scintillation counter. The amount of bound glucose or glyoxal was quantitated and expressed as a molar ratio of glucose or glyoxal per mg BSA.

### 2.4 Serum Albumin Uptake by Bovine Pericardium

Serum albumin uptake kinetics were quantified by measuring the dry and wet mass change of BP samples following BSA exposure. Square samples (n=5, ca. 20mm by 20mm, >18mg dry weight) were cut from glutaraldehyde-fixed BP. The samples were exhaustively rinsed in deionized water to remove any residual salts. Samples were lyophilized for at least 48 hours and their dry weight was measured. The samples were subsequently immersed in 5 ml deionized water per sample for 24 hours gently shaking at 4°C to rehydrate. The wet weight of each sample was measured by blotting each side with Whatman paper (Maidstone, United Kingdom) to remove excess water. The samples were next incubated for 1, 3, 7, 14, or 28 days in solutions (5ml per sample) of PBS with either: BSA (5%), BSA (5%) + glucose (100mM) or BSA (5%) + glyoxal (50mM). All incubations had sodium azide (0.1%) added for sterility. Following incubations, the samples were again exhaustively rinsed with deionized water and lyophilized for 48 hours to calculate dry weight. Wet mass was recalculated as per the previous method.

### 2.5 Morphological Studies

#### 2.5.1 Sample preparation

To prepare samples for immunohistochemistry (IHC) and two-photon microscopy endpoints, samples (n=5) were punched from glutaraldehyde-fixed BP using an 8mm biopsy punch. The BP samples were incubated for 28 days in (1ml per sample) PBS or PBS with either: BSA (5%), glucose (100mM), glucose (100mM) + BSA (5%), glyoxal (50mM), or glyoxal (50mM) + BSA (5%). A group of 5 BP samples were set aside after 24 hours of each incubation to provide a baseline samples for two-photon microscopy. Sodium azide (0.1%) was added to maintain sterility. BP samples were then exhaustively rinsed with PBS prior to follow-up protocols.

#### 2.5.2 Immunohistochemistry Assays

Tissue designated for IHC was fixed in 10% neutral buffered formalin at 4 °C for 48 hours. The tissue was then gradually dehydrated and embedded in paraffin. The paraffin blocks were sectioned at 6μm and mounted on Histobond (VWR; Radnor, PA) slides. The slides were then heated in an oven and re-hydrated in successive xylene to ethanol baths. The slides were then incubated overnight in 60°C citrate buffer (ThermoFisher; Waltham, MA) for antigen retrieval. Following antigen retrieval, slides were rinsed then incubated with primary antibody (αAGE, 0.4 μg/ml | Abcam, Cambridge, United Kingdom; αCML, 0.12 μg/ml | Abcam; αGlucosepane, 4.5μg/ml | David Spiegel laboratory, Yale University, New Haven, CT, per material transfer agreement; αBSA, 0.05 ug/ml| Abcam) overnight at 4°C. Samples to be stained using α-glucosepane were first incubated overnight at room temperature with NaBH_4_ (80mM) in order to reduce glutaraldehyde reaction products that could cross-react with the antibody [23]. Samples were then were incubated with primary antibody as previously described. After incubation with primary antibodies, slides were washed and incubated with H_2_O_2_ (3%) for 10 minutes. Slides were then rinsed and incubated for 1 hour at room temperature with the appropriate horseradish peroxidase polymer-conjugated secondary antibody (Abcam; Cambridge, United Kingdom). Slides were then rinsed and incubated for 8 minutes at room temperature with 3,3'Diaminobenzidine substrate (Abcam; Cambridge, United Kingdom). Slides were then counter-stained using regressive hematoxylin staining, dehydrated, and cover-slipped.

#### 2.5.2 Two-photon Microscopy

All two-photon microscopy scans were performed using a custom Prairie Technologies Ultima Multiphoton Microscopy on Olympus BX-61 upright microscope (Olympus; Tokyo, Japan). This system is equipped with GaAsP photomultiplier tubes and a tunable femtosecond laser (Spectra Physics, MaiTai DeepSee). The system is capable of providing multi-color imaging including second harmonic generation (SHG) imaging. Tissues designated for two-photon microscopy scans were mounted on chamber slides and immersed in PBS. Scans were performed with excitation at 980nm and scanning from the surface to as deep as could be resolved at 5μm steps.

### 2.6 Uni-axial Testing and Related Data Analyses of Collagen Sponges

Dog bone shaped samples (8mm wide that narrows to 4mm wide in the middle, 25mm long) were punched from the glutaraldehyde fixed CS for the mechanical tests. The samples were each incubated for 7 days at 37°C shaking at 110RPM in 5ml per sample of PBS or PBS with either: glucose (100mM), glyoxal (50mM), BSA (5%), glucose (100mM) + BSA (5%), or glyoxal (50mM) + BSA (5%). All incubations contained sodium azide (0.1%) to maintain sterility. Samples were extensively rinsed with PBS and stored in PBS at 4°C until the mechanical tests. Prior to testing, samples’ cross-sectional area was calculated by measuring the cross-section’s length and width using a micrometer. The mechanical testing consisted of: preloading to 0.03N (~3x the sample wet weight), 10 cycles at 1Hz of preconditioning going from 0% to 10% strain, a relaxation test rapidly (14mm/s) extending to 20% strain and holding for 60s, a 2-minute recovery period at 0 strain, then a slow extension (0.05mm/s) to failure. All mechanical testing was performed at the Penn Center for Musculoskeletal Disorders (University of Pennsylvania; Philadelphia, PA) on an Instron® 5542 system (Instron; Norwood, MA). Load (N) and extension (mm) measurements were recorded alongside images of samples following preloading. Samples that did not break cleanly in the center or that slipped during testing were excluded.

The time-load-extension data were analyzed using custom Matlab (Mathworks; Natick, MA) scripts to extract viscoelastic, linear elastic, and failure mechanical properties. Zero strain length was determined by pixel measurement of clamp to clamp distance following preloading. Viscoelastic metrics calculated were the degree of relaxation (the fraction of stress attenuation during relaxation, 1 - σ_equilibrium_/σ_peak_) and time constant of relaxation curve. Linear elastic metrics calculated were the elastic modulus (slope of the stress-strain curve in the linear loading region, expressed as MPa) and the engineering strain at start of the linear loading region of the stress-strain curve. Failure metrics calculated were the ultimate tensile strength (the stress at mechanical failure, expressed as MPa) and the engineering strain at failure. The mechanical data were normalized to the PBS incubation data for each repetition before comparisons.

### 2.7 Statistical Methods

The significance of the effects of BSA presence on 28-day glucose and glyoxal incorporation was determined by two-sample t-test (2.3.1). A two-sample t-test was also used to compare the 28-day incorporation of glucose and glyoxal on fresh and glutaraldehyde fixed bovine pericardium (2.3.2). It was also considered whether the assumptions of the t-test are warranted and the nonparametric analogue, Wilcoxon rank sum, was used instead. To evaluate the significance of the change in dry mass at 28 days, Dunnett’s method was used to compare all BSA groups to the PBS control while Tukey’s HSD test compared each BSA incubation to one another (2.3.3). Dunnett’s method was also used to determine the significance of the changes from the PBS control for each of the metrics from the mechanical tests on the CS (2.6). For all statistical tests, p<0.05 was considered significant. All data are expressed as mean ± standard deviation.

## 3. RESULTS

### 3.1 Model glycation studies

These experiments sought to characterize the kinetics and extent of glucose and glyoxal incorporation into BP and CS in order to model AGE formation in these materials. BP was chosen for use in these experiments because of its use in BHV leaflets, and CS was investigated as a model collagenous material, hypothetically comparable to BP, and better suited for the mechanical studies presented later in this paper. The approach for these studies was to assess the overall potential for AGE formation using both ^14^C-glucose and ^14^C-glyoxal incorporation as exemplary glycation reagents. The rationale for these studies was based on established AGE formation reactions involving glyoxal, a reactive intermediary derived from glucose that specifically reacts with lysine and arginine residues[15]. Furthermore, glyoxal is involved in the formation of carboxy-methyl-lysine (CML), an AGE involved in the pathophysiology of diabetes and other diseases[24–26]. Serum proteins are AGE modified[24,27]; thus, their influence on ^14^C-glucose and ^14^C-glyoxal incorporation was modeled in these studies of BP and CS using serum albumin, the most abundant serum protein, at physiologic concentrations in the specific protocols.

Glucose incorporation, without BSA, was observed to reach 37.4 ± 8.53 nmol/mg in BP at 24 hours (Figure 1A) and, by comparison, in CS the 24-hour incorporation was 73.26 ± 2.27 nmol/mg (Figure 1B). Glucose incorporation leveled off at 7 days, and plateaued by 28 days reaching 295.19 ± 22.84 nmol/mg in BP (Figure 1A) and 367.84 ± 10.99 nmol/mg in CS (Figure 1B). Glucose incorporation in the presence of BSA had comparable kinetics to glucose alone: leveling off at 7 days and plateauing by 28 days in both BP and CS. Co-incubation with BSA resulted in glucose content of 28.96 ± 2.12 nmol/mg in BP (Figure 1A) and 48.23 ± 14.16 nmol/mg in CS at 24 hours (Figure 1B). After 28 days, glucose levels for the co-incubation with BSA reached 103.6 ± 7.26 nmol/mg in BP (Figure 1A) and 124.38 ± 3.68 nmol/mg in CS (Figure 1B).

**Figure 1.**
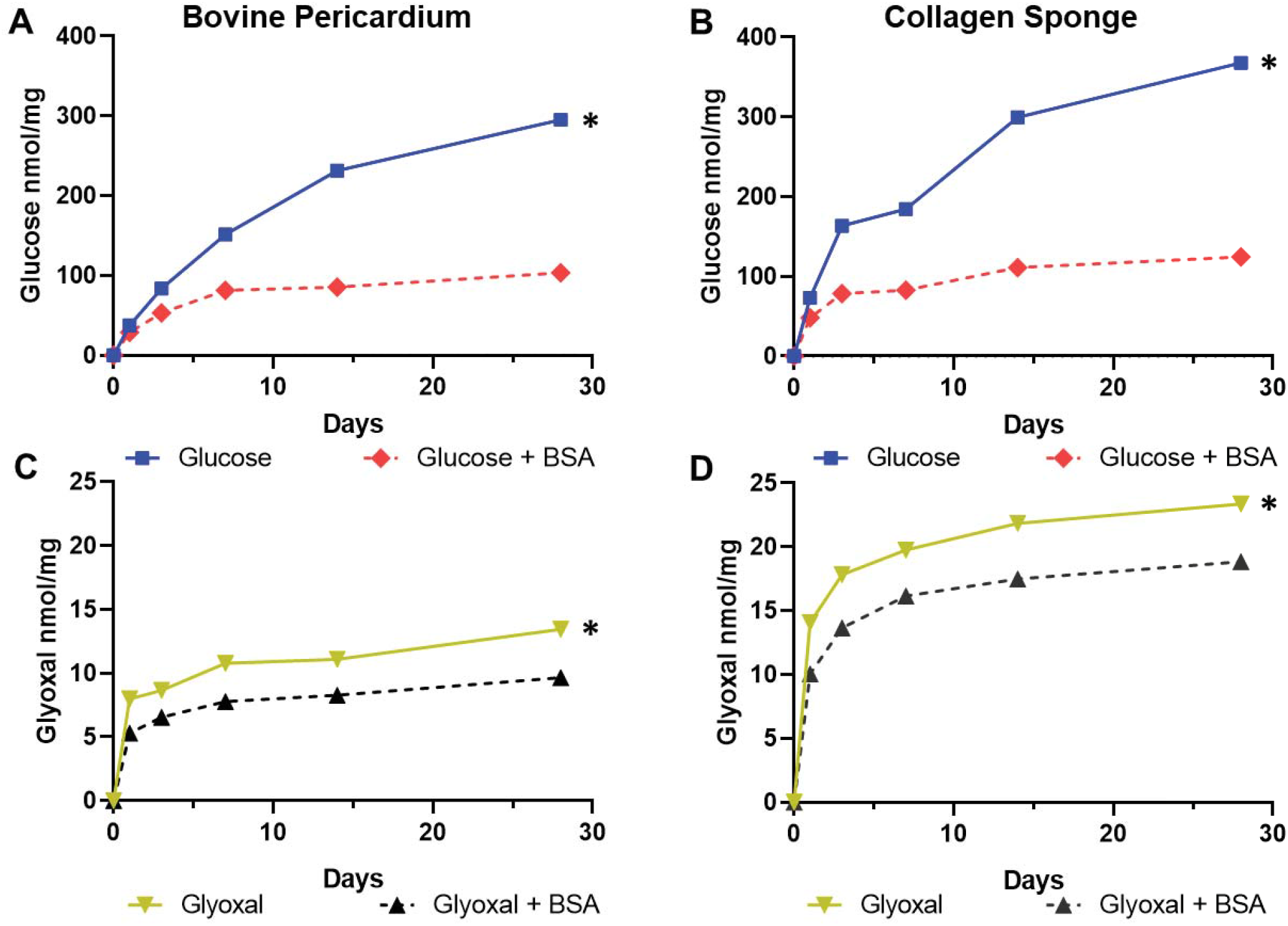
Glycation of glutaraldehyde pretreated bovine pericardium (BP) compared to glutaraldehyde pretreated collagen sponges (CS): ^14^C-glucose and ^14^C-glyoxal incorporation studies with and without the presence of bovine serum albumin (BSA) to simulate serum protein conditions. A) ^14^C-glucose (100mM) uptake by BP with and without the presence of BSA (5%). * p<0.001 B) ^14^C-glucose (100mM) uptake by CS with and without the presence of BSA (5%). * p<0.001 C) ^14^C-glyoxal (50mM) uptake by BP with and without the presence of BSA (5%). * p<0.001 D) ^14^C-glyoxal (50mM) uptake by CS with and without the presence of BSA (5%). * p <0.001 Data shown are means of 10 replicates for 1A and 1C and means of 5 replicates for 1B and 1D. The standard deviations of these values are too small to show graphically. Significance was determined by two sample t-tests.

Glyoxal incorporation at 24 hours was 7.98 ± 1.54 nmol/mg in BP (Figure 1C) and, by comparison, the 24-hour incorporation in CS was 14.07 ± 0.26 nmol/mg (Figure 1D). Glyoxal incorporation leveled off after 7 days and plateaued at 28 days in both BP and CS. Glyoxal incorporation at 28 days reached 13.45 ± 1.77 nmol/mg in BP (Figure 1A) and 23.36 ± 0.47 nmol/mg in CS (Figure 1D). Co-incubation with BSA resulted in 5.29 ± 0.56 nmol/mg in BP (Figure 1C), and 10.04 ± 0.15 nmol/mg in CS at 24 hours (Figure 1D). Glyoxal incorporation in the presence of BSA appeared to also level off by 7 days and plateau at 28 days in both BP and CS. The plateau levels reached at 28 days were 9.66 ± 0.75 nmol/mg in BP (Figure 1C) and 18.84 ± 0.31 nmol/mg in CS (Figure 1D).

### 3.2 Glycation of bovine pericardium occurs regardless of glutaraldehyde pretreatment

Glutaraldehyde crosslinking is the universally used pre-treatment step for preparing bioprosthetic heart valves. However, the effects of glutaraldehyde pretreatment on AGE formation have not been previously studied. Glutaraldehyde reacts with primary amines, principally lysine, to form Schiff bases and heterocyclic crosslinks. This could hypothetically block glycation sites arising from lysyl amine reactions. To assess the effect glutaraldehyde-crosslinking has on glycation capacity, experiments were performed to compare ^14^C-glucose and ^14^C-glyoxal incorporation into fresh BP and glutaraldehyde-fixed BP.

Fresh BP 28-day glucose incorporation was 304.14 ± 37.02 nmol/mg (Figure 2A). This was similar to the glutaraldehyde-fixed BP glucose incorporation of 281.85 ± 10.27 nmol/mg by day 28. In the co-incubation with BSA, glucose incorporation at 28 days was 100.86 ± 13.80 nmol/mg into the fresh BP and 107.51 ± 8.12 nmol/mg into the glutaraldehyde-fixed BP (Figure 2A). By 28 days, the glyoxal incorporation into the fresh BP was 13.25 ± 0.10 nmol/mg (Figure 2B). The glyoxal incorporation into the glutaraldehyde-fixed BP was 14.59 ± 0.90 nmol/mg. In the glyoxal-BSA co-incubation, the 28-day glyoxal incorporation was 8.80 ± 0.23 nmol/mg into the fresh BP and 10.14 ± 0.30 nmol/mg into the glutaraldehyde-fixed BP (Figure 2B).

**Figure 2.**
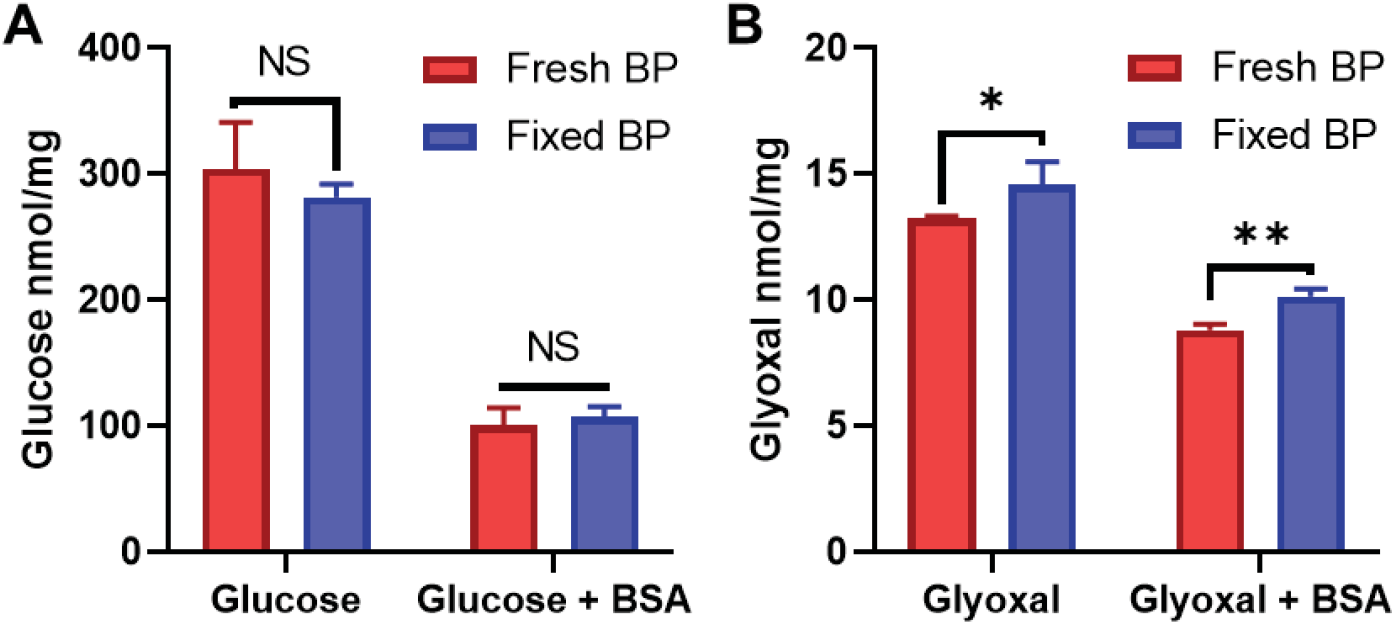
Glycation of non-crosslinked bovine pericardium (BP) compared to glycation of glutaraldehyde-crosslinked BP: ^14^C-glucose and ^14^C-glyoxal incorporation studies. A) ^14^C-glucose (100mM) incorporation into fresh or glutaraldehyde pretreated BP, with or without co-incubation in BSA (5%), was studied as above (Figure 1). No significant differences in incorporation due to glutaraldehyde pretreatment were noted (NS, not statistically significant). B) ^14^C-glyoxal (50mM) incorporation with or without BSA (5%) into fresh BP versus glutaraldehyde-fixed BP was also studied. Glutaraldehyde pretreated BP demonstrated a small but significant increase in glyoxal incorporation (* p=0.028, ** p<0.001). Data shown are 5 replicates. Error bars indicate standard deviation. Significance was determined by two sample t-tests.

### 3.3 Model studies of BSA glycation

BSA co-incubations significantly altered the glycation levels of BP and CS (Figure 1); therefore, the following studies investigated the glycation kinetics of BSA. BSA was incubated in the presence of ^14^C-glucose or ^14^C-glyoxal with the resulting incorporation measured. Unlike the BP and CS glycation kinetics (Figure 1), the glycation of the BSA did not plateau by 28 days (Figure 3); both glucose and the glyoxal incorporation into BSA demonstrated comparable kinetics with increasing incorporation over the 28-day time course. Glucose incorporation into BSA was 45.45 ± 3.59 nmol/mg at 24 hours and 72.78 ± 9.49 nmol/mg at 28 days (Figure 3). Glyoxal incorporation into BSA was 8.24 ± 0.84 nmol/mg at 24 hours and 33.38 ± 0.33 nmol/mg at 28 days (Figure 3). Based on the concentrations of glucose and glyoxal, 3.64 ± 0.47% of the glucose and 3.33 ± 0.03% of the glyoxal are bound to the BSA by day 28.

**Figure 3.**
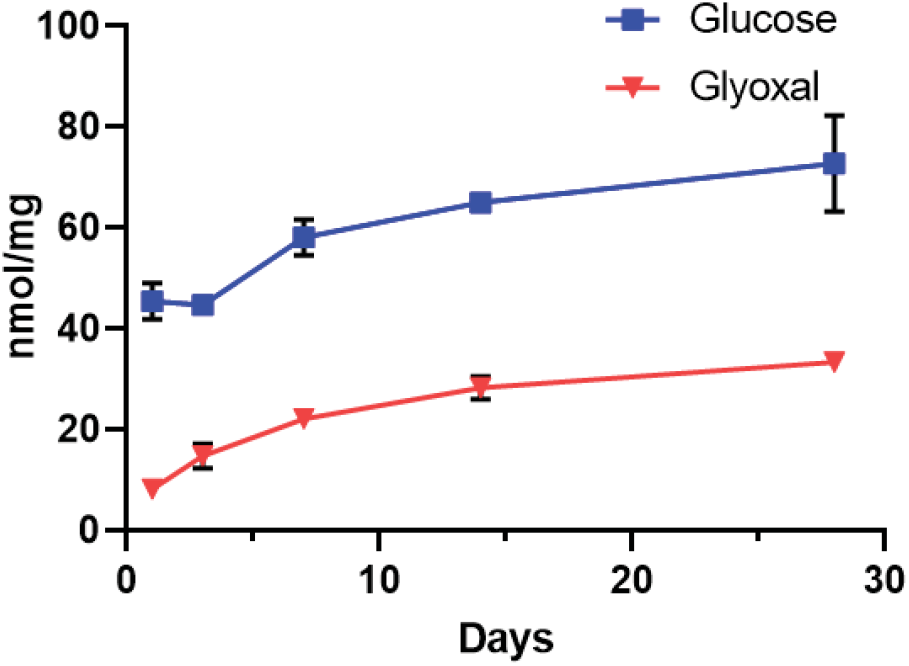
The kinetics of glycation of bovine serum albumin: ^14^C-glucose and ^14^C-glyoxal incorporation studies. The 28-day time courses of ^14^C-glucose (100mM) or ^14^C-glyoxal (50mM) incorporation into 5% bovine serum albumin are shown. Data shown are 3 replicates. Error bars indicate standard deviations.

### 3.4 Bovine serum albumin mass uptake by bovine pericardium

BP samples were incubated in BSA, BSA with glucose, and BSA with glyoxal to model BHV serum protein exposure under different glycation conditions with the resulting BP mass change quantitated. Albumin addition was quantified as the percent change in the dry (Figure 4) and wet weight (not shown) of BP samples from the dry and wet weight before a 1-to 28-day incubation. The dry weight data showed a 4.00 ± 1.15% cumulative loss of mass in the PBS incubation by day 28 (Figure 4). Incubation in BSA offset this loss of dry weight, causing a net increase of 1.84 ± 0.90% by day 28 (Figure 4). Glucose presence in the glucose-BSA co-incubation demonstrated no significant effect on the dry mass change relative to BSA by itself with a 28-day mass increase of 1.06 ± 0.91% (Figure 4). By contrast, glyoxal presence in the glyoxal-BSA co-incubation significantly increased the dry weight gain relative to both the PBS incubation and the BSA-only incubation, reaching a net increase of 6.24 ± 1.85% after 28 days (Figure 4). The change in wet weight data was inconsistent, with too great a variance for statistically significant differences between treatments to be observed.

**Figure 4.**
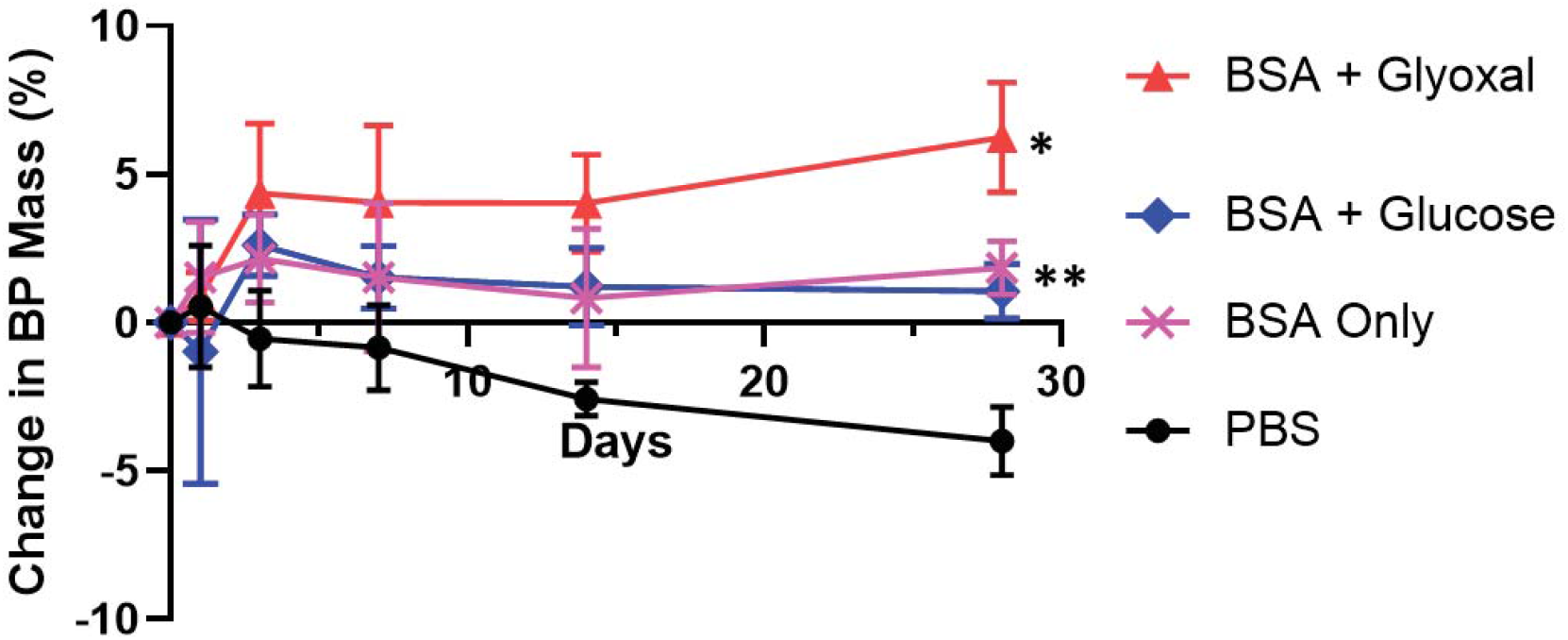
The changes in mass of glutaraldehyde-crosslinked bovine pericardial samples (BP) using glycation conditions in the presence of bovine serum albumin (BSA) to simulate serum protein exposure. Change in dry mass of BP samples over 28-day time course incubating in either: PBS (control), BSA (5%), BSA (5%) with glucose (100mM), or 5% BSA (5%) with glyoxal (50mM). Data shown are percentages of the change in weight from the starting measures with 5 replicates for each condition. Error bars indicate standard deviation. *Comparison by Dunnett’s method showed BSA + glyoxal had significantly different mass change from PBS (p<0.001). Comparison using Tukey’s HSD showed significant differences between BSA + glyoxal and both BSA (p<0.001) and BSA + glucose (p<0.001) at 28 days. ** Comparison by Dunnett’s method showed BSA and BSA + glucose each had significant changes relative to PBS at 28 days (p<0.001). Comparison using Tukey’s HSD test showed no significant difference between BSA and BSA + glucose at 28 days.

### 3.5 Protein glycation as demonstrated by immunostaining of bovine pericardium

These studies sought to characterize the morphologic distribution of protein glycation resulting from the incubation conditions described above. Glycation precursors, such as glucose and glyoxal, have a large family of intermediaries, including Amadori products, on the pathway to forming the mostly irreversible AGE[15]. The antibodies used in these experiments were specific for: CML, an AGE derived from glyoxal[15,24]; general AGE formation; and glucosepane, the most common physiological crosslinking AGE[28]. The BP samples were incubated for 28 days to correspond with the radioactive assays (Figure 1) and two-photon microscopy scans (Figure 6). The CS samples were incubated for 7 days to correspond with the mechanical tests (Table 1).

An array of representative micrographs of immunohistochemistry-stained incubated BP following a 28-day incubation are shown in Figure 5. The α-CML antibody lead to moderate staining in the glyoxal incubated sample (Figure 5E) and heavy staining in the glyoxal-BSA co-incubated sample (Figure 5F) relative to the PBS control (Figure 5A). The α-AGE antibody prompted increased staining only in the glucose incubated sample (Figure 5I) relative to the PBS control (Figure 5G). Likewise, the α-glucosepane antibody demonstrated increased staining over the PBS control only in the glucose incubated samples (Figure 5O vs 5M). The samples exposed a BSA incubation (Figure 5T, 5V and 5X) each showed significant staining with the α-BSA antibody relative to the PBS control (Figure 5S) with the darkest staining occurring in samples from the glyoxal-BSA co-incubation (Figure 5X).

**Figure 5.**
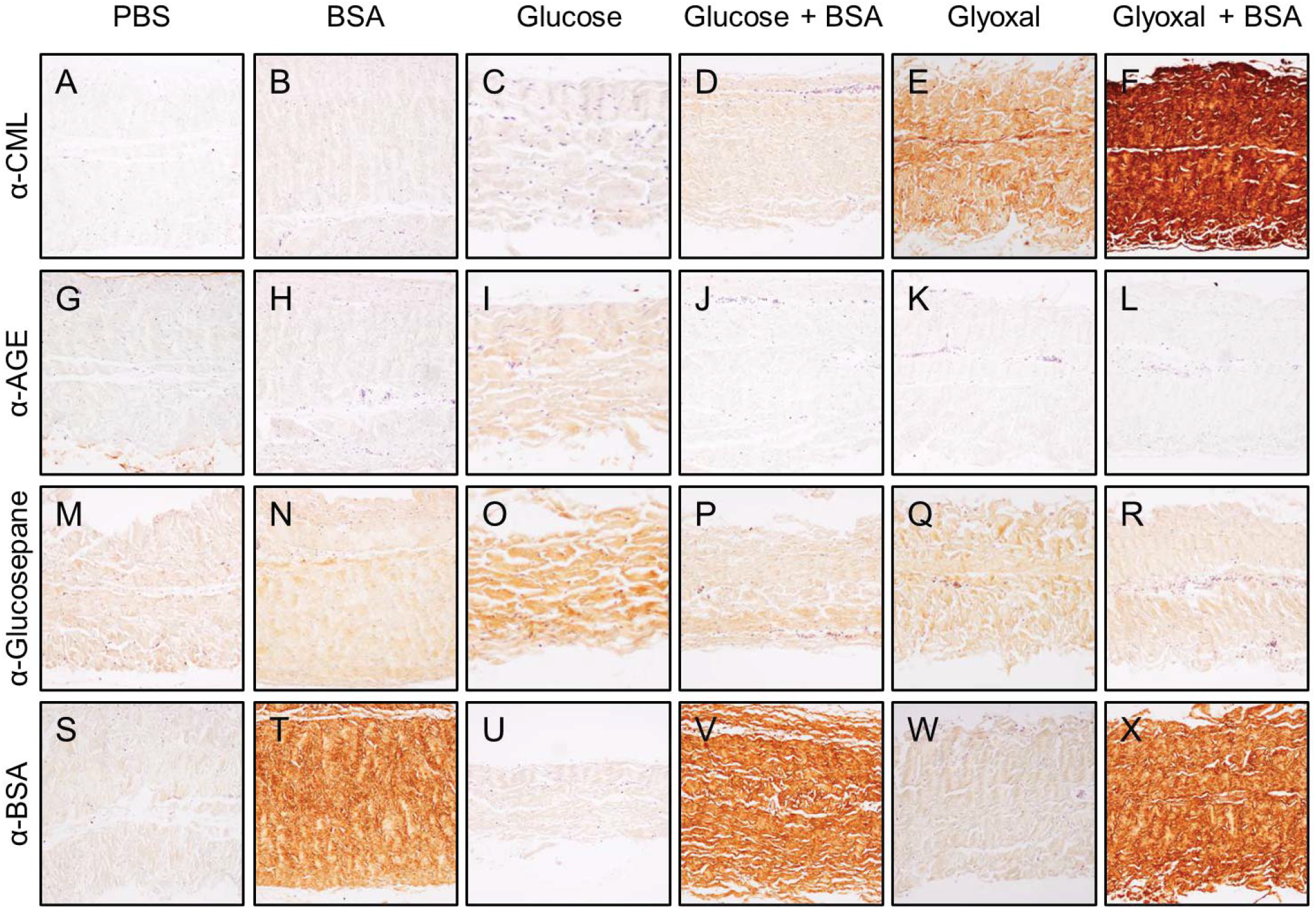
Formation of advanced glycation end products (AGE) in glutaraldehyde-crosslinked bovine pericardium following *in vitro* incubation: glycation-specific immunohistochemistry micrographs of glutaraldehyde-crosslinked bovine pericardium incubated for 28 days using the conditions indicated: A-F) Staining for carboxy-methyl-lysine (CML); G-L) Staining for general AGE formation. M-R); Staining for glucosepane formation. S-X); Staining for bovine serum albumin (BSA). Immunoperoxidase staining was used with a substrate of 3,3'Diaminobenzidine; Incubation concentrations were: bovine serum albumin (BSA, 5%), glucose (100mM), and glyoxal (50mM). Original magnification 100x.

**Figure 6.**
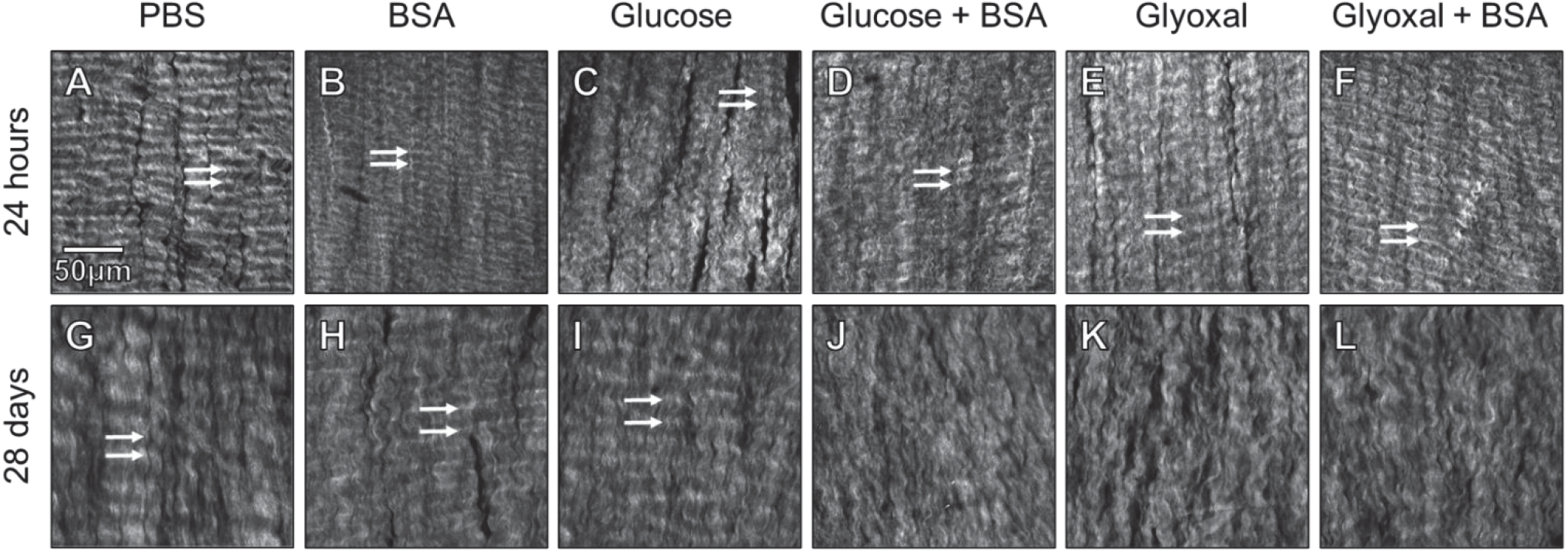
Two-photon microscopy images of glutaraldehyde-fixed bovine pericardium following *in vitro* incubations (see Methods): Samples were incubated for 24 hours (A-F) or 28 days (G-L) in the following media: A&G) PBS, B&H) bovine serum albumin (BSA, 5%), C&I) glucose (100mM), D&J) glucose (100mM) plus BSA (5%), E&K) glyoxal (50mM), and F&L) glyoxal (50mM) plus BSA (5%). All images are presented at the same scale. Arrows indicate typical crimp spacing where bands are discrete.

### 3.6 The effects of protein glycation on BP collagen structure—two photon microscopy results

To study the effects of glycation and serum protein exposure on BHV’s collagen microarchitecture, two-photon microscopy scans were performed on BP samples following *in vitro* incubations. BP samples after 24 hours of incubating consistently demonstrated collagen fibers were aligned with a tight crimp period and distinct crimping bands under all conditions (Figure 6A-F). Incubating BP in PBS for 4 weeks caused some alterations in the structure, but no significant loss of alignment nor a change in crimp period relative to 24 hours of PBS incubation (Figure 6G). After 4 weeks of BSA exposure, there is a significant loss of identifiable crimp and an increase in the crimp period relative to the image at 24 hours and 4 weeks of PBS, however, the fibers retain the bulk of the orientation and still have distinct crimp bands (Figure 6H). Glucose exposure, by itself, had relatively mild effects that were similar to BSA exposure, some loss of crimp, a significant increase in crimp period relative to the 24 hours scans and 4 weeks of PBS exposure, but the BP retained collagen alignment and banding (Figure 6I). However, glucose-BSA co-incubation produced a dramatic disruption in the collagen microarchitecture, almost completely eliminating any crimp bands with significant collagen misalignment (Figure 6J). After 4 weeks of glyoxal exposure, the fibers are significantly maligned and the crimp bands are completely lost (Figure 6K). The glyoxal-BSA co-incubation produced the greatest modification of the structure after 4 weeks with compete loss of any crimp banding (Figure 6L).

### 3.7 The effects of glycation on the viscoelastic properties of collagen sponges

To evaluate the effects glycation and serum proteins have on glutaraldehyde-fixed collagen’s mechanics, uni-axial mechanical tests were performed on CS samples following 7-day *in vitro* incubation. CS was selected as there was no dominant directionality to the collagen fibers to complicate uni-axial tests. Furthermore, CS was found to have highly consistent and uniform mechanical and chemical properties in pilot studies. The 7-day duration was selected as glucose and glyoxal incorporation into the CS leveled off at 7 days (Figure 1B & 1D). The mechanical tests consisted of a relaxation test, followed by a recovery period, then a slow extension to failure.

The effects of each treatment on the viscoelastic, linear elastic, and failure mechanical properties of the collagen sponges are expressed as changes from the PBS incubation (Figure 7). Glucose both by itself caused a significant loss in relaxation (8.09 ± 7.90%, Figure 7A) with no significant changes in any other metric. Glyoxal exposure by itself produced a significant loss of relaxation (14.51 ± 4.08%, Figure 7A), a significant decrease in the strain to reach failure (8.20 ± 10.51%, Figure 7B), a significant increase in stiffness (10.48 ± 13.47%, Figure 7C). BSA by itself caused significant increase in elastic modulus (10.57 ± 15.42%, Figure 7C) and increase in ultimate tensile strength (8.58 ± 8.69%, Figure 7D). The glucose-BSA co-incubation was similar to glucose alone with the only significant change being a loss of relaxation (16.63 ± 9.99%, Figure 7A). The glyoxal-BSA co-incubation had effects similar to the BSA-only incubation with a significant increase in stiffness (10.05 ± 18.23%, Figure 7C) and a marginal increase in ultimate tensile strength (4.80 ± 12.58%, p=0.22, Figure 7D) relative to the PBS control.

**Figure 7.**
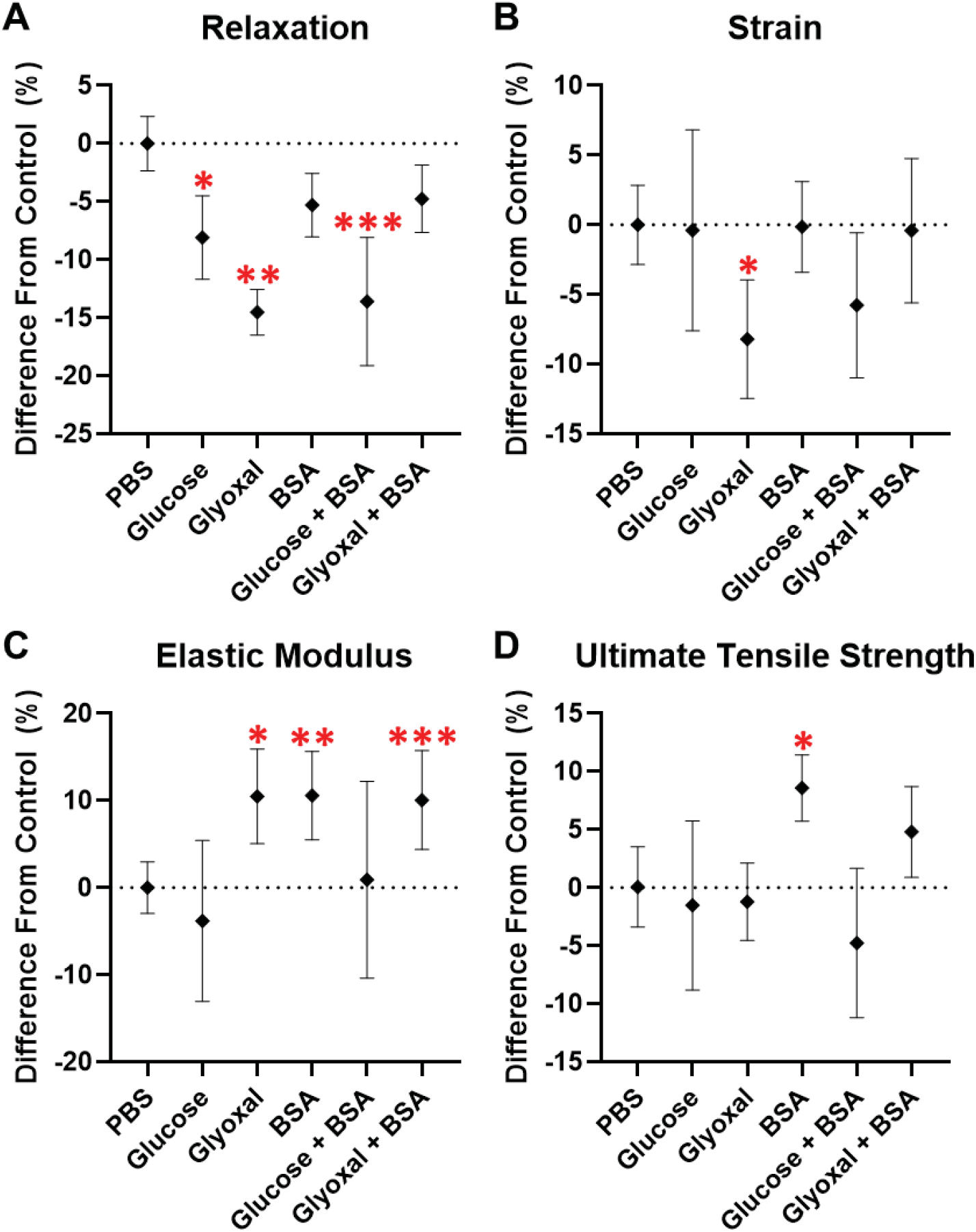
Changes of mechanical properties determined with uniaxial testing of collagen sponge samples following 7-day incubations. Results are normalized relative to phosphate buffered saline (PBS) controls. A) Degree of relaxation, relative stress attenuation during relaxation. Significant decreases were observed with glucose (*p=0.0057), glyoxal (** p<0.001), and glucose plus BSA (*** p<0.001). B) Strain at mechanical failure. Significant decrease was observed with glyoxal (* p=0.026). C) Elastic modulus. Significant increases were observed with glyoxal (* p=0.030), BSA (** p=0.010), and glyoxal plus BSA (*** p=0.012). D) Ultimate tensile strength. Significant increase was observed with glyoxal (* p=0.004). Incubation conditions were: PBS (control) [n=46], glucose (100mM) [n=21], glyoxal (50mM) [n=19], bovine serum albumin (BSA, 5%) [n=38], glucose (100mM) plus BSA (5%) [n=16], and glyoxal (50mM) plus BSA (5%) [n=36]. Error bars indicate 95% confidence interval. Significance was determined by Dunnett’s method comparing to PBS.

## 4. DISCUSSION

The present studies provide novel information concerning the susceptibility of BHV to glycation-related pathophysiology. Our results documented BHV glycation kinetics, processes by which glycation contributes to BHV SVD, and the deleterious effects of serum proteins that contribute to BHV glycation. These model system studies demonstrated that BHV are highly susceptible to rapid glycation (Figure 1) by both glucose and glyoxal, with or without serum albumin, and produced IHC results in BP *in vitro* comparable to failed clinical explants (Figure 5) [23]. Furthermore, the present studies show that AGE formation in BP occurs at sites independent of those involved in glutaraldehyde crosslinking, since fresh BP demonstrated comparable glucose and glyoxal incorporation as glutaraldehyde pretreated BP (Figure 2). These model studies used SHG to demonstrate disruption of collagen fibril organization (Figure 6) that was virtually identical to that seen in failed clinical explants[23]. Glucose effects modeled *in vitro* in the present studies, were not examined in our original report. However, glucose had a far greater level of incorporation than glyoxal (Figure 1A versus Figure 1C), and disrupted collagen structure (Figure 6) of BP with a morphology comparable to clinical explants[23]. It was also observed that there was significantly reduced glucose and glyoxal incorporation into BP and CS for the BSA co-incubations (Figures 1&2). The mechanisms responsible for this are complex, and the diminished glucose and glyoxal uptake were most likely due to BSA reactively binding to sites that would otherwise have reacted with glucose or glyoxal. BSA has a multitude of amino acids capable of engaging in the formation of AGE, including lysines, arginines and histidines. These BSA glycation-reactive amino acids are also involved in the formation of complex heterocyclic AGE as well as AGE-related crosslinks. Taken together this provides the basis for understanding our observation of the differential incorporation of glucose and glyoxal into BP and CS in the presence of BSA.

CS, as a model for the BP collagen network, made possible uniaxial mechanical testing that documented alteration of the viscoelastic and linear elastic properties in CS due to AGE formation and serum protein incorporation (Figure 7). This means that through a cardiac cycle, the instantaneous load on glycated BHV tissue would be greater and the material would have less capacity to disperse the stress. These mechanical results provide a mechanistic explanation for our previous in vitro pulse duplicator findings that demonstrated AGE formation caused hydrodynamic dysfunction of clinical grade tricuspid BHV that resulted in decreased effective orifice area and increased pressure gradient[23]. These dysfunctional effects are directly related to clinical SVD. The present results suggest that the hydrodynamic changes observed in the pulse duplicator studies are likely due to increased stiffness and loss of viscoelasticity making the valve leaflets more resistant to cyclic opening.

The present results concerning the reaction kinetics of glucose and glyoxal with BP (Figure 1A&C), while not previously reported in BHV studies, are comparable to previous model system results by others using radiolabeled glycation reagents [29–33]. For example, carboxymethyl lysine formation from ^14^C-glyoxal in bovine serum albumin has been shown in studies by others to have comparable reaction kinetics to those observed in our BHV investigations[30]. In addition, prior studies of ^14^C-glucose incorporation into retinal basement membranes[33] demonstrated reaction rate results that were also comparable to those for BP in our experiments. These comparisons indicate that the susceptibility of BHV to glycation occurs through mechanistic pathways that are operative in other pathophysiologies.

SHG analyses provided novel insights concerning collagen structural alterations in BP due to glycation and serum albumin uptake, such as collagen malalignment loss of collagen crimp (Figure 6). Collagen crimp, that structurally represents collagen’s folding during its unloaded state, is important to BHV function and its loss causes is indicative of diminished compliance and increased susceptibility to mechanical injury[34]. This study’s collagen modifications were comparable to the changes observed using SHG in our clinical study evaluating failed clinical BHV tissue, *in vivo* rat subdermal explanted BP, and *in vitro* pulse duplicator experiments[23]. Taken together these observations validate our current *in vitro* models as reproducing representative alterations in collagen structure, comparable to clinical explant BHV leaflets with SVD [23].

There are some caveats and limitations of this study to be noted. The concentrations of reagents used in the glycation studies were adjusted to provide an accelerated model system. An isotonic 5% BSA solution was used. However, glyoxal was utilized at 50mM, based on prior model studies using this reagent [23,30,35,36], rather than the physiologic concentration (~154nM)[37]. Glucose, at 100mM, was also used at diabetic level, hyperglycemic concentrations rather than physiologic (~5.6mM)[38] levels. Nevertheless, this glucose concentration was representative of concentrations used in previous accelerated *in vitro* glycation studies by others[31–33,39–41]. Furthermore, glucose was not investigated in our prior AGE-BHV study of clinical explants[23], and is of strong relevance because of the more rapid SVD BHV failure process in diabetics[14]. Thus, the glucose results in the present results addressed the important contributions of glucose mediated glycation of BHV to SVD. The IHC assays of the study focused on well known, representative immunostaining targets reported in other glycation studies and these were: AGE, carboxymethyl lysine and glucosepane. These same IHC markers were endpoints in our clinical study[23] as well as other clinical studies[25,42–47], and thus have mechanistic and clinical relevance. Uniaxial testing of CS showed that glycation results in loss of viscoelasticity and increased stiffness (Figure 7). BP was not studied because of the well-known technical challenges of the anisotropic nature of BP[48–50]. As discussed earlier, the present uniaxial testing observations account for the degeneration of the hydrodynamic properties of BHV following glycation and serum protein modifications observed in our pulse duplicator study[23].

The results of this study have several implications addressing SVD. AGE-mitigating strategies for use with BHV have not been previously investigated; the presents results indicate anti-AGE agents may be of interest. Currently available anti-AGE treatments include AGE inhibitors and AGE breakers. AGE inhibitors are pharmacologic agents, e.g. aminoguanidine and pyridoxamine, that prevent the formation of AGE by mechanisms such as scavenging the reactive carbonyl intermediates and blocking the oxidation of the Amadori intermediate, respectively[51]. AGE breakers, e.g. phenacyl-thiazolium bromide and alagebrium, function by cleaving glycation crosslinks and have been demonstrated to have the capacity to prevent or reverse glycation modifications in vascular disease[52,53]. The present study’s model systems are capable of evaluating in vitro those treatments’ efficacy at preventing or reversing the glycation modifications of BHV. Furthermore, our studies demonstrated the impact of serum protein infiltration in vitro on BHV structure and mechanical properties, and provided insights to explain clinical and experimental observations. Lastly, diabetic patients are at higher risk for earlier SVD[14,54,55]. Since the present studies showed significant effects of glucose on structure and mechanical properties of BP and CS, future *in vivo* model studies using diabetic animals represent an important direction. A notable example of such a study investigated both collagen and elastin scaffolds implanted subdermally in control and diabetic rats[56]. The samples explanted from diabetic rats demonstrated significantly increased biaxial stiffness compared to controls.

## 5. CONCLUSIONS

The results of the present study support the hypothesis that glycation and serum protein infiltration can contribute to SVD pathophysiology and lead to the following conclusions: 1) BP and CS are susceptible to rapidly progressive glycation and serum albumin incorporation; 2) BSA, with our without glyoxal or glucose, disrupts collagen structure; 3) Glucose alone, while not dramatically altering collagen structure after 28 days compared to PBS, does alter viscoelastic properties; 4) Similarly, collagen’s uniaxial properties are significantly altered by both glycation and serum albumin incorporation. Taken together glycation and serum protein infiltration contribute to SVD via the structural changes in collagen that were observed in these studies and the related disruption of viscoelastic and linear elastic properties.

## Authorship contribution statement

**Christopher A. Rock:** Conceptualization, Methodology, Software, Formal analysis, Investigation, Writing - Original Draft, Project administration. **Samuel Keeney:** Methodology, Investigation, Writing - Original Draft, Writing - Review & Editing. **Andrey Zakharchenko:** Methodology, Investigation, Writing - Original Draft. **Hajime Takano:** Methodology, Software, Resources. **David A. Spiegel:** Resources. **Abba M. Krieger:** Formal analysis. **Giovanni Ferrari:** Conceptualization, Resources, Supervision, Funding acquisition. **Robert J. Levy:** Conceptualization, Methodology, Resources, Writing - Review & Editing, Supervision, Funding acquisition.

## Disclosure

Robert J. Levy is a consultant for WL Gore. This does not represent a conflict of interest related to this publication. David A. Spiegel is a shareholder of Revel Pharmaceuticals, and this does not represent a conflict of interest for the present studies. Christopher A. Rock, Samuel Keeney, Andrey Zakharchenko, Hajime Takano, Abba M. Krieger, and Giovanni Ferrari have no competing interests to disclose.

## ACKNOWLEDGEMENTS

Biomechanical testing was performed at the Penn Center for Musculoskeletal Disorders (NIH AR069619). This work was supported by NIH R01s HL122805 (GF) and HL143008 (RJL and GF), T32s HL007915 (RJL and CR), HL007854 (APK) and HL007343 (AF), The Kibel Fund for Aortic Valve Research (to GF and RJL), The Valley Hospital Foundation ‘Marjorie C Bunnel’ charitable fund (to GF and JG), the American Diabetes Association Pathway to Stop Diabetes Grant 1-17-VSN-04 and the SENS Research Foundation (to DAS), and both Erin’s Fund and the William J Rashkind Endowment of the Children’s Hospital of Philadelphia (to RJL).

## TABLE OF ABBREVIATIONS

AGE: advanced glycation endproducts
BHV: bioprosthetic heart valves
BP: bovine pericardium
BSA: bovine serum albumin
CML: carboxy-methyl-lysine
CS: collagen sponge
IHC: immunohistochemistry
PBS: phosphate buffered saline
SHG: second harmonic generation
SVD: structural valve degeneration

## *Declaration of Interest Statement

### Rock et al: Declaration of Interests Statement

The following competing interests are disclosed: Robert J. Levy is a consultant for WL Gore. This does not represent a conflict of interest related to this publication. David A. Spiegel is a shareholder of Revel Pharmaceuticals. However, this does not constitute a conflict of interest concerning the present manuscript and its contents. Christopher A. Rock, Samuel Keeney, Andrey Zakharchenko, Hajime Takano, Abba M. Krieger, and Giovanni Ferrari have no competing interests to disclose.

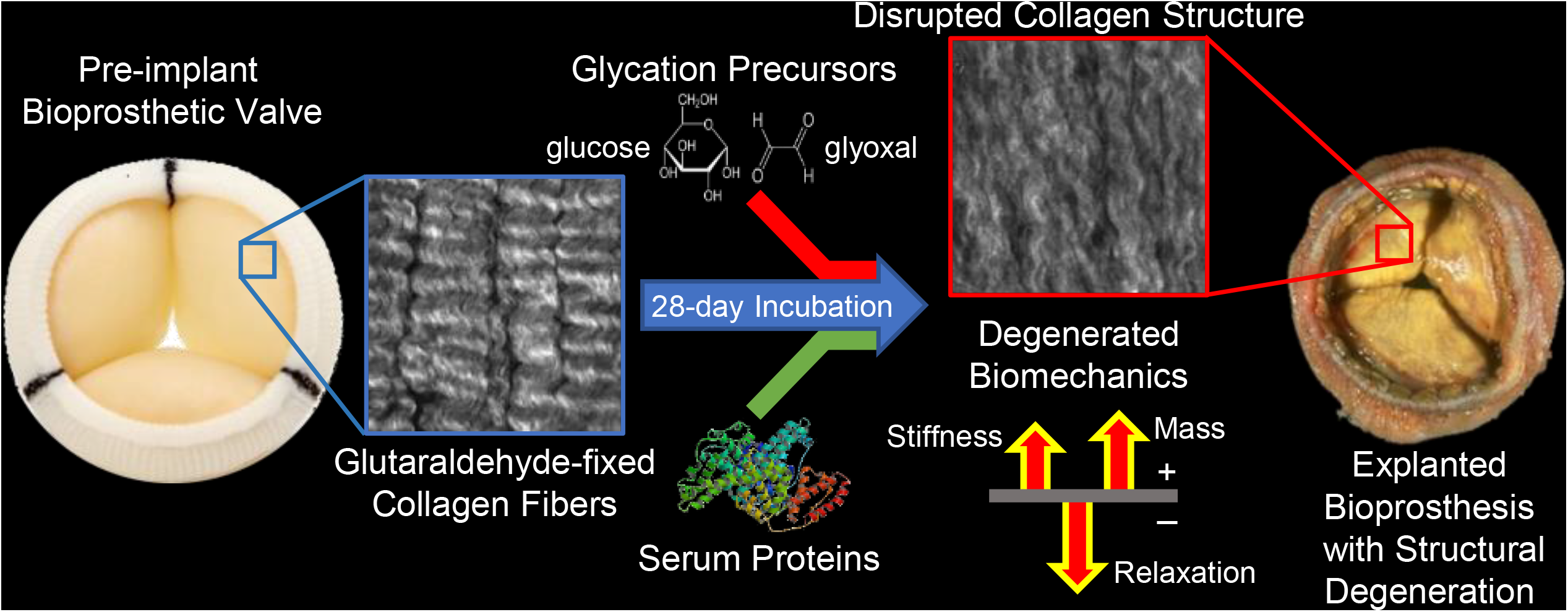

